# Cutting epitopes to survive: the case of lambda variant

**DOI:** 10.1101/2021.08.14.456353

**Authors:** Stefano Pascarella, Massimo Ciccozzi, Martina Bianchi, Domenico Benvenuto, Roberto Cauda, Antonio Cassone

**Author notes:** Equal contribution (Stefano Pascarella, Massimo Ciccozzi). Corresponding author, Prof. Massimo Ciccozzi, Medical Statistic and Molecular Epidemiology Unit, University of Biomedical Campus, 00128 Rome, Italy.

## Abstract

This manuscript concisely reports an in-silico study on the potential impact of the Spike protein mutations on immuno-escape ability of SARS-CoV-2 lambda variant. Biophysical and bioinformatics data suggest that a combination of shortening immunogenic epitope loops and generation of potential N-glycosylation sites may be a viable adaptation strategy potentially allowing this emerging viral variant escaping host immunity.

## Introduction

A number of SARS-CoV-2 variants termed variants of interest (VOI) or concern (VOC) have attracted investigators’ attention to elucidate the mechanisms underlying their enhanced transmission and/or resistance to neutralization by immune sera and monoclonal antibodies [1,2]. Among more recently identified VOI, the lambda variant, which has emerged in Peru and has rapidly spread to South American regions and US remains poorly investigated, particularly regarding the effects of mutations on the thermodynamic parameters affecting the stability of the Spike protein and its Receptor Binding Domain (RBD). Variations in these parameters, if consistent, have been shown to predict relevant changes in SARS-CoV-2 contagiousness and immunoescape ability [3,4].

## Materials and methods

For this study, we retrieved the Spike sequence representative of the lambda variant from the GISAID [5] translated protein set of the isolate coded as EPI_ISL_1629764. The reference Wuhan Spike sequence is labelled by the RefSeq [6] code yp_009724390. We modelled the mutations in the Spike Receptor Binding Domain (RBD) onto the crystal structure denoted by the PDB code 6M0J using the functions available within the graphic program UCSF Chimera [7].The Spike deletion has been modelled using the method available in the web server SwissModel [8] using as a starting structure the coordinate set identified by the PDB id 7KRS. This structure has been selected by SwissModel as the best trimeric template. This coordinate set contains the Cryo-EM structure of the Spike mutant D614G. The point mutations have been introduced within the obtained model using the functions incorporated in UCSF Chimera [7]. The servers Dynamut and GlycoPred [9] and NetNGlyc [10] were used to predict point mutations impact onto the Spike structure, and glycosylation sites, respectively. Protein-protein interaction energy has been predicted with the method implemented in PRODIGY [11]. Structural analysis and visualization have been carried out with PyMOL [12] or UCSF Chimera [7].

## Results and discussion

We first notice that mutations G75V and T76I occur at an exposed loop connecting two short antiparallel β-strands (**Table 1 and Figure 2**). The effect of each of the two mutations is predicted to be stabilizing. This loop is in contact to the loop encompassed by the sequence positions 246-280 that is one of the epitopes recognized by mAbs [13]. Interestingly, the deletion 246-252 (corresponding to the sequence RSYLTPG) occurs within this loop. To predict the potential impact of the deletion onto the NTD affinity to a human mAbs, the complex between SARS-CoV-2 Spike and 4A8 Ab deposited as 7C2L in PDB has been used as a case study. The mutant NTD has been modelled via SwissModel and superposed to the wild-type domain of the complex. Interface interactions and energies calculated by PRODIGY have been compared (**Table 2 and Figure 2**). Overall, the partial deletion of the loop is predicted to weaken interaction to the 4A8 antibody with consequent decrease of binding affinity as several interactions are removed. In particular, deletion in lambda NTD removes interactions that in the wild-type complex take place between L249 and F60, Y54 of the 4A8 light chain. Moreover, a salt-bridge and a π-cation interaction between R246 and 4A8 E31 and Y27 respectively disappear in the lambda variant (**Figure 2**).

**Figure 1.**
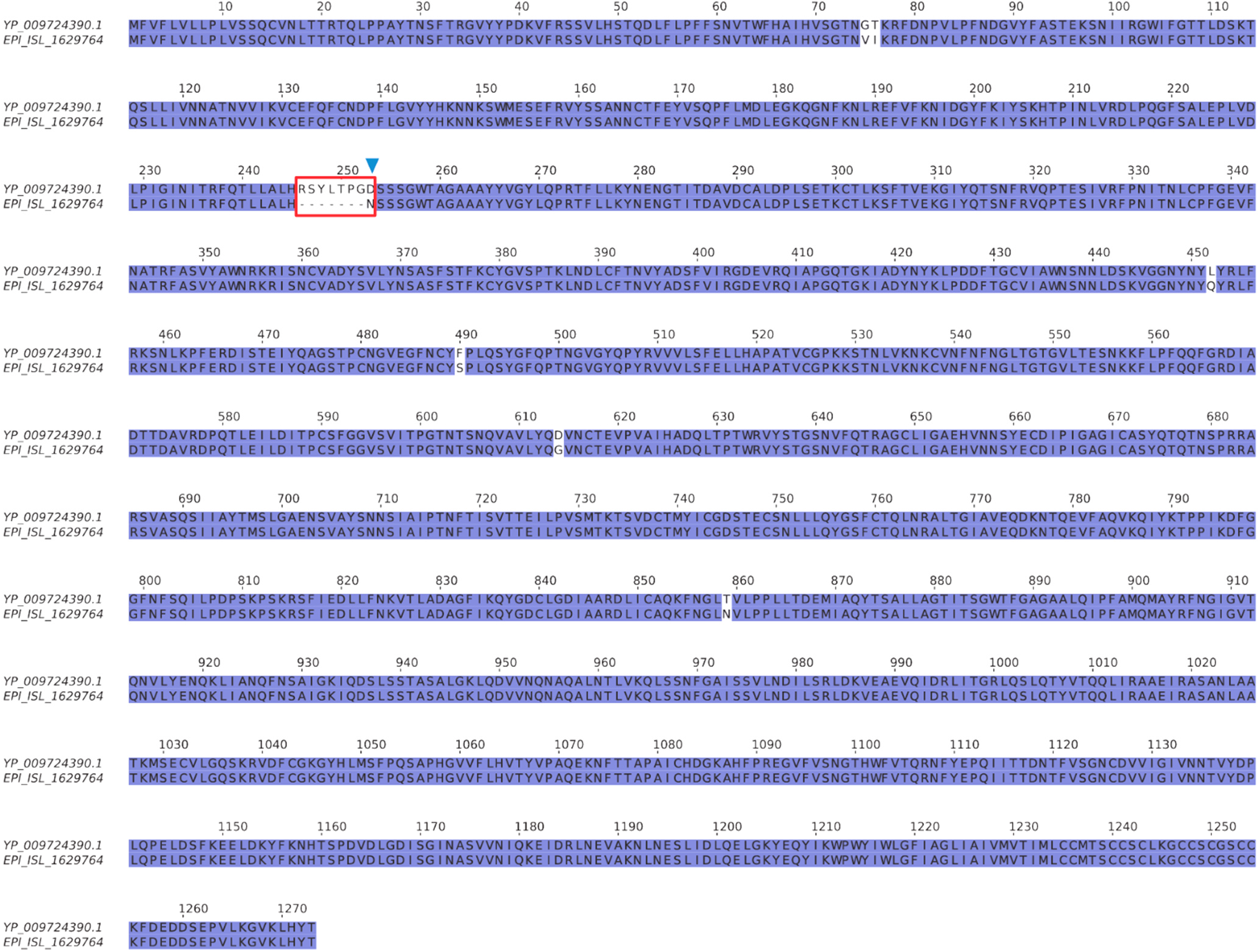
Sequence alignment between the Wuhan reference (yp_009724390.1) and the lambda variant (EPI_ISL_1629764) Spikes. Red box indicates the deletion of the NTD loop while the green triangle marks the potential new N-glycosylation site.

**Figure 2.**
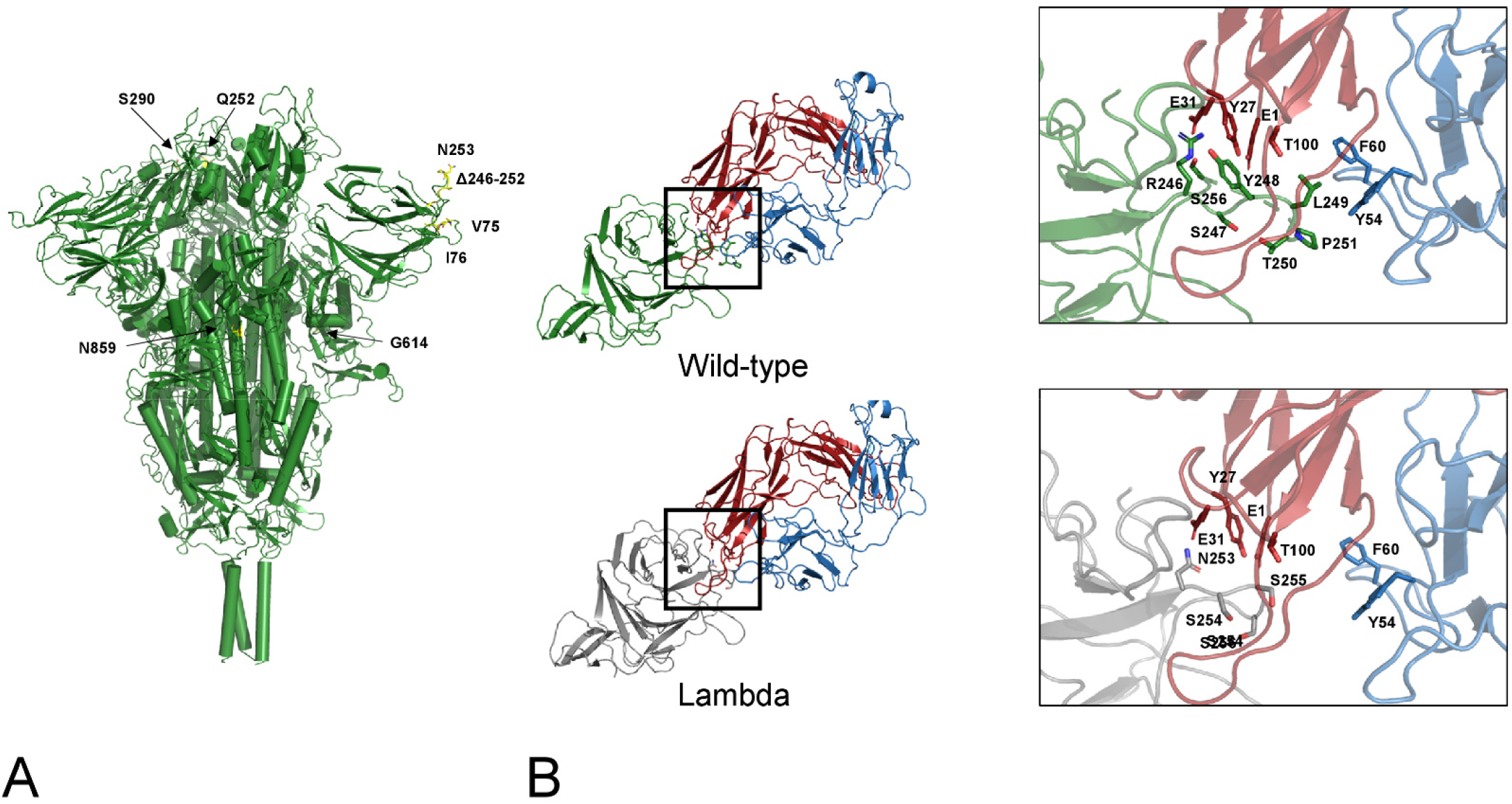
A) Map of the lambda mutations onto the trimeric Spike structure. Only one mutation per monomer has been indicated. B) interface NTD-4A8. The mutant NTD has been modelled via SwissModel and superposed to the wild-type domain of the complex. Interface interactions and energies calculated by PRODIGY have been compared. Comparison of the interfaces between the wild-type (green cartoon) and the lambda NTD (grey cartoon) and the mAb 48A at the deletion region. Red and blue cartoons indicate the heavy and light chain respectively. Side chains in the deleted loop and the interacting residues are indicated with stick models and labelled. Sequence numbering refers to the wild-type Spike.

**Table 1.** Characterizing mutations of the lambda variant Spike protein with respect to the reference Wuhan sequence.

| Mutation | Structural position | $\Delta\Delta G$ (kcal/mol)<sup>a</sup> | Local interactions |
| --- | --- | --- | --- |
| G75V | NTD. β-hairpin. Exposed | 0.214 (moderately stabilizing) | Hydrophobic interaction to V57 |
| T76I | NTD. β-hairpin. Buried | 1.248 (highly stabilizing) | Hydrophobic interaction to L244, W251. Adjacent to the deleted loop 246-252 |
| $\Delta 246-252$ | NTD. Loop | Not applicable<sup>b</sup> | Deleted loop involved in interaction with human antibodies (4A8, FC05) |
| D253N | NTD deletion C-terminal side | Not applicable<sup>b</sup> | Introduces a potential N-glycosylation site |
| L452Q | Receptor binding motif | 0.295 (moderately stabilizing) | Increases surface hydrophilicity. Close to the interface to ACE2 |
| F490S | Receptor binding motif | -0.039 (neutral) | Increases surface hydrophilicity. Possible polar interaction to E484. Close to the interface to ACE2 |
| D614G | SD2 | Not tested in this work | Extensively characterized. Influences conformational flexibility and susceptibility to proteolytic activation. |
| T859N | S2. Short β-strand | -0.037 (neutral) | H-bond to K847 of the same chain. In proximity of G614 of the other chain |

a) Results from Dynamut.
b) Dynamut does not process deletions or simultaneous mutations

**Table 2.**
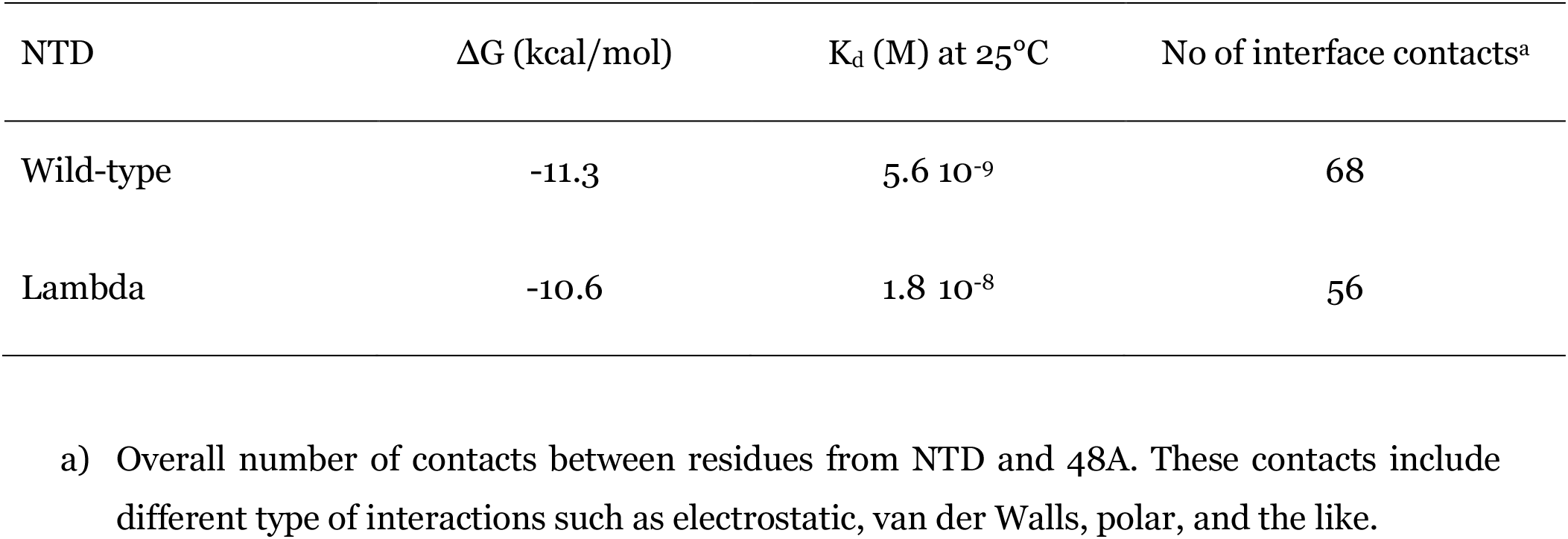
PRODIGY prediction between wild-type reference and lambda Spike NTD interaction energies to 4A8 Ab.

In contrast to the above, the mutations in the RBD region (L452Q and F400S) are predicted to have little impact on the domain thermodynamic stability (**Table 1**). Both replacements change a hydrophobic residue into a hydrophilic one inducing a local increase of surface hydrophilicity. The sites are exposed to the solvent close to the interface RBD-ACE2. The binding affinity of the mutant and wild-type RBD is almost identical according to the PRODIGY predictions (results not shown). Finally, of the two the mutations within the S1 and S2, the worldwide abundant and conserved D614 G has been amply described and attributed with transmissibility increase (….). The T859N occurs in the S2 part of the Spike in a short β-strand in a position that is about 8 Å distant from G614 of the adjacent subunit. The impact of the mutation onto the thermodynamic structure is predicted to be neglectable (**Table 1**). However, this mutation introduces a H-bond to K847. It cannot be excluded that this mutation in cooperation with D614G can result in a synergic long-distance allosteric effect on Spike conformational flexibility. Indeed, allosteric effects of mutations in this portion of the Spike have been suggested and studied [14,15].

Overall, our in-silico analysis suggests that the point mutations characterizing the lambda variant do not seem to directly influence RBD affinity for ACE2 receptor, thus making it uncertain a relevant impact on virus transmission, unless the mutations in the S2 domain have a long-distance allosteric effect on conformational flexibility and ACE2 affinity. The most evident and likely functionally impacting change of the lambda variant is represented by the 246-252 deletion since they could confer to the virus an enhancing capacity to escape the host immune response by two theoretic though likely and already reported strategies; shortening epitopes located in the loops and exploiting increased glycosylation. variations in spike cell epitopes and glycosylation profile during virus transmission have been already described [16].

While this manuscript was being prepared for submission, we came across a pre-print published in BioRxiv by Izumi Kimura and collaborators (bioRxiv doi: https://doi.org/10.1101/2021.07.28.454085;) where the N-terminus 7 amino acids deletion is described and discussed for its potential relevance in conferring to the lambda variant of SARS-CoV-2 resistance to antiviral immunity. Though with different approach, the data by Kimura and collaborators are in total agreement with our own from in silico methods.

